# Tumor cell specific total mRNA expression informed neural networks predicts cancer progression

**DOI:** 10.64898/2026.05.01.722212

**Authors:** Ankita Paul, Jessica C. Lal, Shuangxi Ji, Carissa Fong, Kevin Chen, Yu Ding, Ruonan Li, Yaoyi Dai, Quang Tran, Matthew Montierth, Saverio Alberti, Scott Kopetz, Wenyi Wang

## Abstract

Inferring tumor molecular phenotypes from high-dimensional multi-omic data is a fundamental challenge in computational biology. Current methods for estimating tumor cell–specific total mRNA expression (TmS) require matched DNA and RNA sequencing data and rely on computationally intensive deconvolution pipelines. We present TmSNet, a deep learning framework that predicts TmS using mRNA, DNA methylation, miRNA, and immune cell proportions as input features. TmSNet integrates structured feature selection (gradient boosting, LASSO, elastic net) with specialized neural architectures to predict continuous TmS. Across 12 TCGA cancer types, TmSNet achieved cross-validated performance up to concordance correlation coefficient (CCC) = 0.93 and correlation R² = 0.88 and generalized to external cohorts with correlations of 0.54 (SCAN-B) and 0.43 (FUSCC). Predicted TmS values effectively stratify patients by risk and preserve known transcriptional profiles across tumor subtypes. These results demonstrate that TmSNet can infer biologically meaningful phenotypes from multi-omic data and provide a scalable framework for modeling tumor transcriptional activity in heterogeneous cohorts.

## Introduction

Cancer progression and treatment resistance are a result of dynamic transcriptional programs that emerge from clonal evolution and microenvironment selection^1–3^. Yet precision oncology remains disproportionately focused on driver gene mutations, largely overlooking tumor-intrinsic transcriptional changes that shape treatment response and disease prognosis^4–6^. Although genomic characterization of cancers has enabled the development of targeted therapies against oncogenic drivers, producing strong but often short-lived responses, mutation-centric frameworks fail to capture the system-level transcriptional heterogeneity of evolving tumors^7,8^. Importantly, emerging evidence suggests global tumor gene expression, not just individual gene expression, has provided insights to aneuploidy and myc-activation, reflecting biological programs that support tumor growth^9–11^. However, there remains a critical gap in scalable approaches to quantify these global transcriptional phenotypes in cancer patient cohorts. Our prior work demonstrated that variation in total tumor mRNA expression (TmS) is a robust prognostic marker across 15 cancer types^12,13^. TmS integrates transcriptomic and genomic deconvolution, accounting for tumor transcript proportion, purity, and ploidy to isolate tumor-specific expression signals. However, direct computation of TmS requires matched bulk DNA and RNA sequencing data, which are often unavailable in clinical cohorts. Even when this data exists, TmS calculation depends on computationally intensive pipelines that are sensitive to noise and batch effects, limiting scalability and reproducibility across datasets. As a result, the practical application of TmS as a biomarker remains constrained to a subset of well-characterized cohorts, highlighting the need for a scalable framework to infer TmS from widely available multi-omics data.

Deep learning (DL) has become a powerful approach for modeling complex biological systems, particularly in genomics and transcriptomics, by capturing non-linear and hierarchical patterns in high-dimensional space that conventional statistical methods often miss^14–17^. However, the application of DL to predict continuous biomarkers, such as TmS, remains relatively unexplored. Recent DL frameworks have aimed at integrating high–dimensional multi-omics data to predict continuous clinical variables or cluster patients. Frameworks such as Flexynesis have implemented diverse neural architectures, to regress continuous clinical variables such as drug sensitivity and patient age directly from bulk molecular profiles^18^. Despite these advances, a pervasive generalization gap remains when such models are applied to external, previously unseen patient cohorts. This failure is frequently linked to dataset domain shift and technical variability across institutions, where absolute numerical regression outputs diverge substantially due to differences in sequencing depth, library preparation, or platform-specific biases^19–22^. Consequently, even models achieving high internal Pearson correlations often exhibit severe calibration drift, rendering their absolute predictions unreliable for establishing clinical dosing thresholds or informing decision–critical interventions in new environments^23–25^. Another major methodological limitation is the existing methods’ focus on collapse continuous regression outcomes into binary or multi–class endpoints, in particular to subgroup patients^26–31^. Although earlier frameworks such as iClusterPlus demonstrated the value of joint latent variable modeling for capturing continuous biological gradients, many contemporary deep learning studies discretized outcomes into categories such as “sensitive” versus “resistant” to maximize reported AUC values^32^. Emerging evidence indicates that classification–centric models are particularly brittle under external validation, largely because they ignore the underlying molecular continuity that regression architectures are specifically designed to preserve^33,34^. Hence, adopting a regression–first paradigm is essential for accurately modeling intra–tumor heterogeneity and capturing transitional cellular states that underlie tumor aggressiveness. By maintaining continuous representations rather than discretized endpoints, future models will be better positioned to reflect the complexity of tumor evolution in clinical settings^35,36^.

We developed TmSNet, a deep-learning framework that integrates structured feature selection with specialized deep neural network-based predictive models to predict the biologically meaningful and continuous values of tumor-specific total mRNA expression (TmS) directly from easy-to-obtain raw multi-omic profiles (**Fig. 1**). As expected with DL, our framework captures complex non-linear biological interactions that extend beyond traditional linear modeling approaches. Hence, our predicted TmS demonstrates high accuracy when compared with the deconvolved TmS and is significantly associated with clinical outcomes, supporting prognostic relevance. Together, these advances position TmSNet as a scalable approach for integrating heterogeneous molecular data into a single, interpretable, and clinically actionable biomarker.

**Figure 1.**
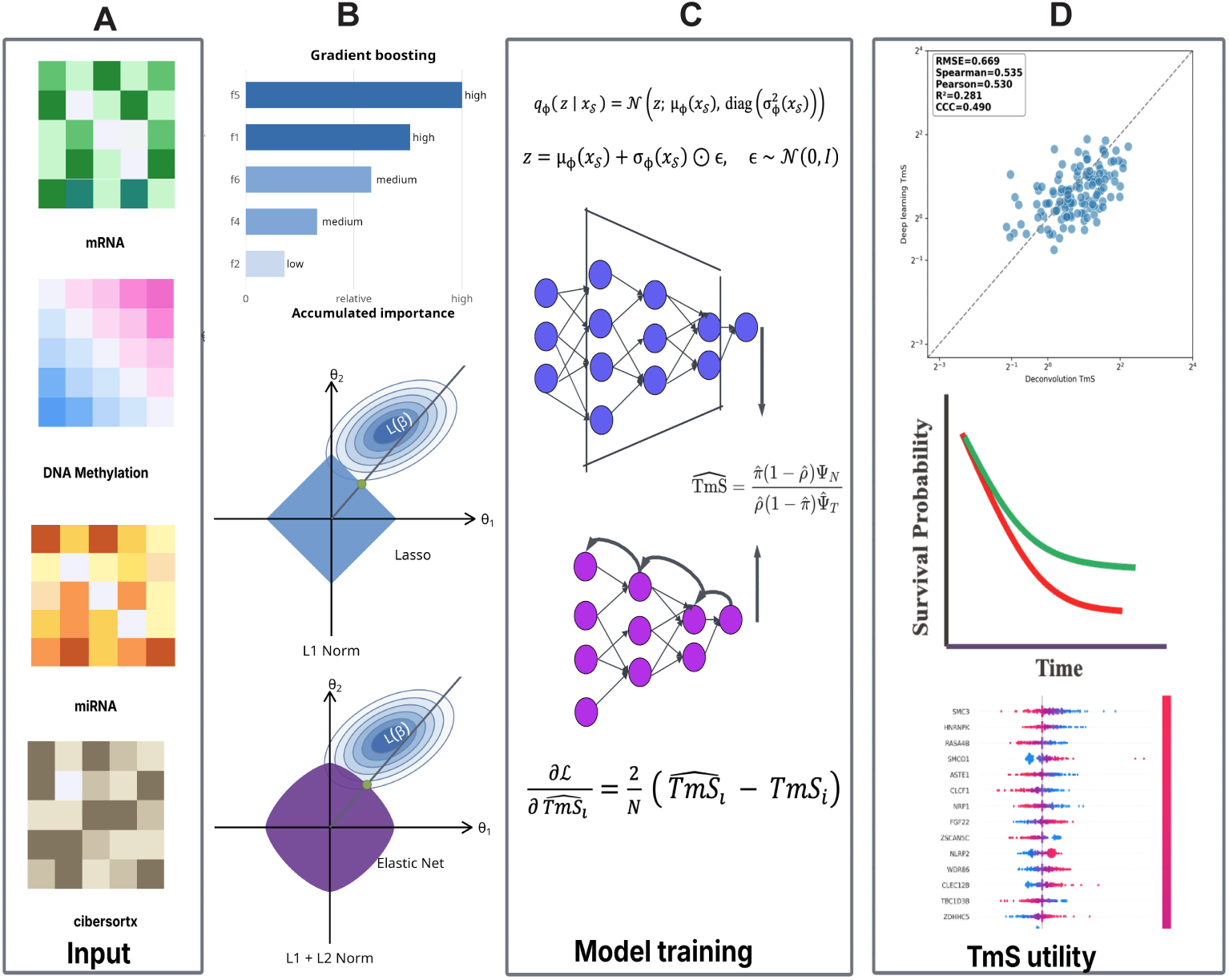
The TmSNet Model Framework. (**A**) Example input -omic profiles, (**B**) Feature extraction and dimensionality reduction using gradient boosting (GB), lasso regression (LR), and elastic net (EN) (**C**) Neural networks training using two types of architecture: specialized encoder neural net (SENN), and feed forward neural net (FFNN) and (**D**) Prognosis validation which includes model performance measured by correlation of predicted total tumor mRNA expression (TmS) and deconvolved TmS, prognostic ability for clinical outcomes, and model explainability.

## Results

### Nonlinear biological dependencies underlying TmS

Our prior work has established TmS as a potential pan-cancer biomarker that is predictive of patient outcomes^12,13^. We showed that high TmS is associated with activation of metabolic programs supporting nucleotide synthesis, including pentose phosphate pathway and glycolysis, as well as transcriptional amplification linked to myc signaling. At the cellular level, elevated TmS reflects stem-like, less differentiated states enriched for epithelial–mesenchymal transition and transcriptional plasticity, independent of proliferation^13^. In specific contexts such as triple-negative breast cancer, TmS additionally captures tumor–microenvironment interactions, distinguishing immune-enriched tumors from stromal/ECM-dominated, immune-excluded states^12^.

We next tested the hypothesis that TmS in breast cancer would be strongly explained by canonical transcriptional regulators or RNA processing machinery. Contrary to expectation, correlations with RNA polymerase subunits, mediator complex components, chromatin remodelers, and splicing factors were weak and inconsistent, with key regulators such as *POLR2A* (Pearson r = −0.38), *MED12* (Pearson r = −0.34), *SF3B1* (Pearson r = −0.35), and *SMARCA2* (Pearson r = −0.49) showing modest associations (**Supplementary Table 1**). Similarly, lineage-defining transcription factors including *FOXA1* (Pearson r = −0.35) and *PBX1* (Pearson r = −0.28) did not positively track with TmS, arguing against a dominant transcription factor–driven model. These findings indicate that TmS cannot be explained by a single-gene regulatory axis, but instead reflects nonlinear, systems-level interactions impacting global transcriptional output that are unlikely to be captured by models single data modality or classic linear models.

In order to capture higher-order relationships that influencing TmS we sought to incorporate multi-omic data. First, we systematically compared baseline machine learning approaches using mRNA alone to deep learning models and multi-omic integration strategies (**Supplementary Table 2**). Across all cancer types, baseline XGBoost models trained on mRNA features showed limited performance, while deep learning architectures applied to mRNA alone provided only modest improvements. In contrast, integrating DNA methylation with mRNA within the TmSNet framework consistently improved predictive accuracy (breast cancer concordance correlation coefficient (CCC): 0.709 to 0.889; bladder CCC: 0.556 to 0.924), indicating that multi-omic inputs capture biological signal not recoverable from transcriptomic data alone. These findings support the need for models that incorporate both nonlinear structure and multiple molecular modalities.

### TmSNet: A Multi-Omic Deep Learning Framework

We developed TmSNet, a deep-learning framework that leverages a modular architecture that integrates advanced machine learning for feature selection with specialized deep neural network-based predictive models to predict tumor-specific total mRNA expression (**Methods**). The input feature space was constructed to capture complementary layers of biological layers influencing global transcription output. Specifically, we included mRNA sequencing, DNA methylation array data, for which the CpG site specific log ratios can be obtained from well-established preprocessing pipelines. Additionally, we included miRNA expression profiles to capture post-transcriptional regulatory signals that complement transcriptional and epigenetic features when modeling global transcript abundance. Immune cell proportions estimated using CIBERSORTx^37^ were included to capture tumor microenvironment heterogeneity. Copy-number variation (CNV) features were excluded because tumor ploidy is directly used in the calculation of TmS and therefore could introduce information leakage and artificially inflate model performance (**Fig. 1A**).

To address the high dimensionality of these inputs, data is first processed through an encoder module incorporating gradient boosting, LASSO regression, and elastic net for dimensionality reduction (**Fig. 1B**). The resulting embedded vectors are used to train specialized deep neural network models variational autoencoders (special encoded neural network or SENN) and multi-layer perceptron (feed forward neural network or FFNN) (**Fig. 1C**). Model outputs are aggregated to generate the predicted total tumor cell specific mRNA score (TmS) (**Fig. 1D**) for downstream interpretation analysis.

First, we performed internal 5-fold cross-validation training from 12 TCGA cancer types (total number of patients = 3,699, cancer-type specific sample size ranges from 155 to 633, **Table 1, Supplementary Fig. 1**). A total of 90 models including 3 feature selections and 15 input data combinations and 2 model designs were tested. The best model in table 1 is chosen by highest R^2^ and CCC and lowest root mean squared error (RMSE). A lower RMSE, a close-to-1 CCC and higher R^2^ correlation would suggest good performance. While the feed forward neural network (FFNN, **Methods**) architecture observes good performance in terms of error rate and correlation to ground truth across different cancers. For most cancers the SENN implementation outperforms it. This performance improvement can be attributed to the latent representation of the non-linear high dimensional realization of the feature vectors that help capture their interactions better to predict TmS. Other than prostate and thyroid cancer, all 10 other cancer types showed good performances with CCC ranging from 0.72 to 0.93. Larger sample size is correlated with the better performance, as expected. DNA methylation data is selected for all cancer types to be predictive of TmS. We further compared the predicted and deconvolved TmS within each fold in each cancer type in order to evaluate stability and robustness in prediction (**Supplementary Fig. 1**) Several cancer types (e.g., breast, colorectal, lung) showed tight clustering near the identity line with uniformly favorable metrics across all five folds, suggesting that the model captures stable signal components of TmS and is not overfitting to partitions. In contrast, cancer types with fewer samples, lower variance, or skewed TmS distributions (e.g., thyroid, prostate) exhibit greater fold–to–fold variability consistent with the known sensitivity of correlation-and error–based measures to dynamic range. The cross-fold stability demonstrates that it is not driven by any single split; rather, our model prediction is sustained across independent test folds, with variability patterns that reflect underlying biological and sampling differences among tumor types. **TmSNet Explainability** To evaluate the relative importance of input features to TmS prediction, we used SHAP (SHapley Additive exPlanations)^38^ values to quantify the contribution of each feature to individual predictions (**Fig. 2**). SHAP values are additive, meaning the sum of feature contributions equals the difference between the model prediction and the baseline prediction for a given sample. We chose five top performing cancer types in the TCGA cross-validation step (**Table 1**) to establish model explainability. In lung, bladder, breast, kidney and pancreatic cancers, top predictive genes displayed a mixture of positive and negative associations with TmS, spanning proliferation, chromosomal instability, extracellular matrix remodeling, immune signaling, differentiation loss, and metabolic rewiring (**Supplemental Figs. 2-4**). In breast cancer, features with the highest weights included *OVCH2* (Pearson r = 0.009, p = 0.82), *ZNF177* (Pearson r = -0.25, p = 6.8 x 10^-10^), *LCLAT1* (Pearson r = 0.04, p = 0.32), *RNF186* (Pearson r = 0.093, p = 0.021) and regulatory factors like *PHF2* (Pearson = -0.52, p < 2.2 x 10^-16^). Multi-omic features also contributed to model predictions, including miRNA MIMAT0000280 and cg14216068, highlighting the added value of multi-omic input features. Most top predictors showed weak or asymmetric correlations with TmS. Collectively, these findings demonstrate that TmSNet captures high-dimensional, non-linear relationships in which the contribution of a given gene depends on context and co-occurring features, rather than on simple additive linear effects.

**Table 1.** Cross validation performance metrics (Mean and STD) across all cancers in the TCGA cohort along with the best feature combination, feature extractor and model training for each cancer type. The model with the best CCC is reported.

| Cancer Type | CV RMSE<br>Mean (STD) | CV CCC<br>Mean<br>(STD) | CV R<sup>2</sup> Mean<br>(STD) | Best Model | Best Feat.<br>Extractor | Best Feat.<br>Combination |
| --- | --- | --- | --- | --- | --- | --- |
| Bladder urothelial carcinoma (n=319) | 0.55 (0.09) | 0.92 (0.03) | 0.87 (0.05) | SENN | ElasticNet | mRNA+Methylation+CIBERSORTx |
| Head and neck squamous cell carcinoma (n=429) | 0.61 (0.05) | 0.93 (0.02) | 0.88 (0.04) | SENN | ElasticNet | mRNA+Methylation+miRNA |
| Prostate adenocarcinoma (n=252) | 0.80 (0.11) | 0.57 (0.09) | 0.41 (0.13) | FFNN | ElasticNet | mRNA+Methylation |
| Kidney renal papillary cell carcinoma (n=155) | 0.67 (0.07) | 0.86 (0.06) | 0.8 (0.08) | SENN | ElasticNet | mRNA+Methylation |
| Kidney renal clear cell carcinoma (n=214) | 0.53 (0.06) | 0.88 (0.03) | 0.80 (0.05) | SENN | Lasso | mRNA+Methylation |
| Colorectal cancer (n=325) | 0.57 (0.06) | 0.79 (0.04) | 0.7 (0.05) | SENN | ElasticNet | mRNA+Methylation |
| Lung squamous cell carcinoma (n=319) | 0.59 (0.11) | 0.72 (0.11) | 0.55 (0.16) | SENN | XGBoost | mRNA+Methylation |
| Lung adenocarcinoma (n=345) | 0.60 (0.06) | 0.81 (0.05) | 0.70 (0.08) | FFNN | ElasticNet | mRNA+Methylation |
| Thyroid carcinoma (n=196) | 0.80 (0.06) | 0.61 (0.04) | 0.44 (0.09) | FFNN | ElasticNet | mRNA+Methylation+miRNA+CIBERSORTx |
| Breast cancer (n=633) | 0.61 (0.04) | 0.89 (0.02) | 0.81 (0.03) | SENN | XGBoost | mRNA+Methylation+miRNA+CIBERSORTx |
| Stomach adenocarcinoma (n=223) | 0.66 (0.13) | 0.89 (0.03) | 0.82 (0.05) | SENN | ElasticNet | mRNA+Methylation+miRNA |
| Uterine corpus endometrial carcinoma (n=289) | 0.70 (0.10) | 0.82 (0.02) | 0.7 (0.03) | SENN | ElasticNet | mRNA+Methylation+miRNA |

**Figure 2.**
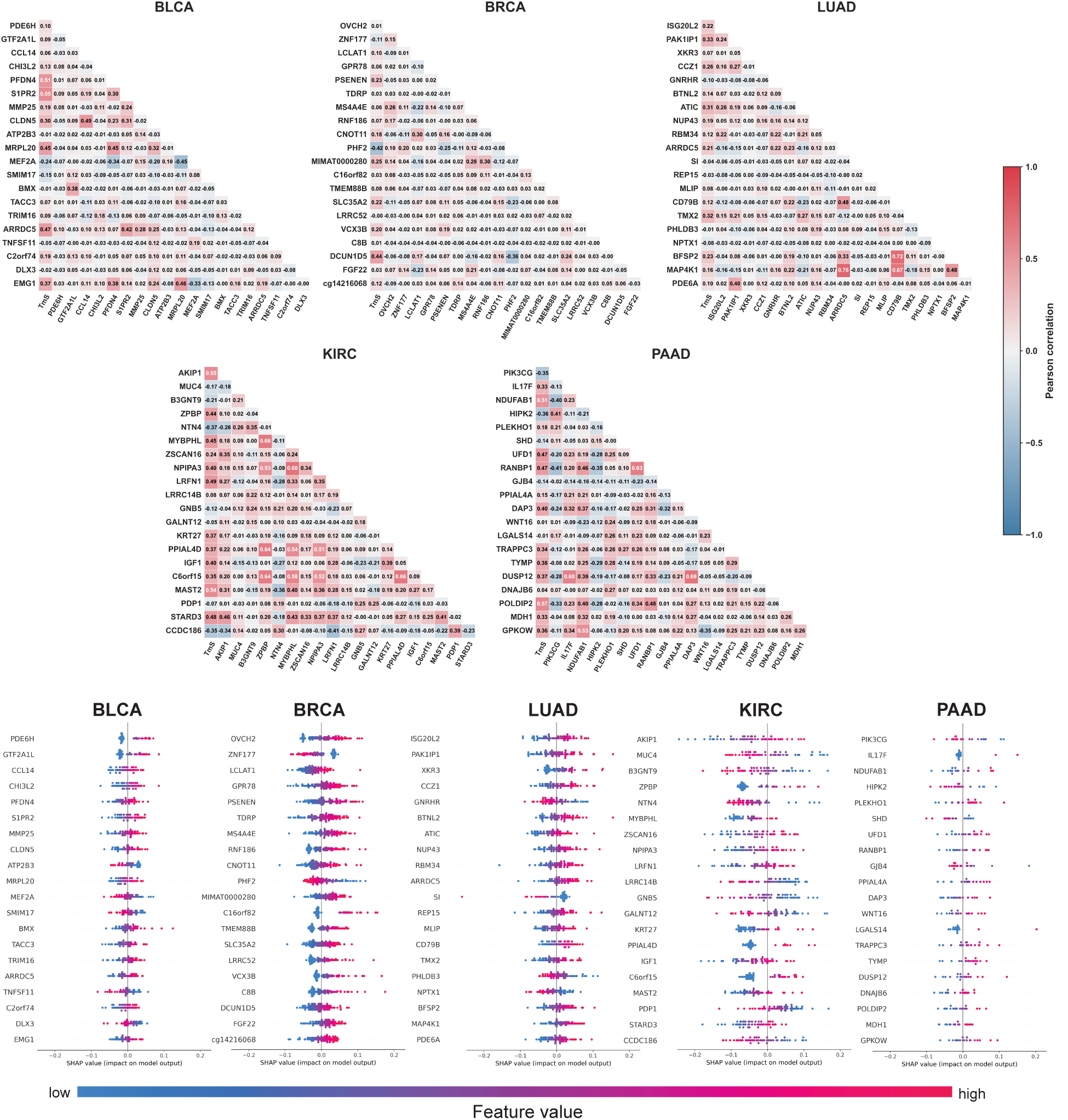
TmSNet explainability for top five cancer types of TCGA cohort with the best RMSE and CCC. Shown are the expression correlations of top 20 features and with TmS, and SHAP values distribution for these features in bladder (BLCA), breast (BRCA), lung (LUAD), renal (KIRC), and pancreatic (PAAD) cancers.

### External Validation

Using TmSNet oversampled and trained from the TCGA breast, we predicted TmS in two unseen cohorts: Fudan University Shanghai Cancer Center (FUSCC)^39^ and the Sweden Cancerome Analysis Network-Breast (SCAN-B)^40^ (**Table 2**). While the utility of TmS has been established in our prior work on TCGA pan-cancer genomic, therapeutic and hazard risk subtyping, our subsequent study in triple-negative breast cancer (TNBC, the most aggressive subtype of breast cancer) observed successful stratification of TNBC subtypes, hazard risk and survival risk prediction across TNBC cohorts^12,13^. This is clinically important as patients with TNBC are still in urgent need of biomarkers to predict responsiveness to chemotherapy and direction towards alternative therapy if not fit for chemo^12^. Hence, here we explore how the predicted TmS from TmSNet drives similar or identical biological interpretations as the existing observations in these cohorts (**Methods**). Due to a lack of DNA methylation data in both cohorts, we expect the predicted TmS be of moderate correlation with the deconvolved TmS.

**Table 2.** External Validation Performance. Shown is a summary of TmSNet performance and survival associations for predicted TmS in external cohorts reporting RMSE, Spearman correlation coefficients, and the predicted-TmS cutoff used to dichotomize tumors.

| Cohort | RMSE | Correlation Coeff. | Continuous Pred. TmS (log2) Cox p | Hazard Ratio (95% CI) | TmS cutoff | Binary TmS LogRank p | Binary TmS Cox p | Hazard Ratio | Continuous TmS Cox p | Hazard Ratio |
| --- | --- | --- | --- | --- | --- | --- | --- | --- | --- | --- |
| SCANB (131) | 0.67 | 0.54 | 0.1 | 0.63 (0.32–1.25) | 1.54 | 0.1 | 0.15 | 0.63 (0.33–1.19) | 0.16 | 0.69 (0.41–1.16) |
| FUSCC (218) | 1.6 | 0.43 | 0.3 | 0.76 (0.44–1.32) | 1.94 | 0.03 | 0.07 | 0.57 (0.31–1.05) | 0.2 | 0.78 (0.52–1.15) |

Across the external cohorts, the violins and scatterplots together show that TmSNet captures the overall ordering of TmS but with cohort-specific interpretation. In SCAN-B, the predicted distribution broadly matches the true TmS (similar spread and median) (**Fig. 3A-B**), and the test-set metrics indicate moderate agreement (RMSE = 0.669; Spearman r = 0.535; Pearson r = 0.530; R² = 0.281; CCC = 0.490), with points clustering around the identity line and only mild under/overestimation at the extremes. In FUSCC, the predicted values are markedly compressed toward the lower range consistent with range restriction in the scatter plot, producing metrics (RMSE = 1.608; Spearman r = 0.431; Pearson r = 0.416; R² = 0.173; CCC = 0.249).

**Figure 3.**
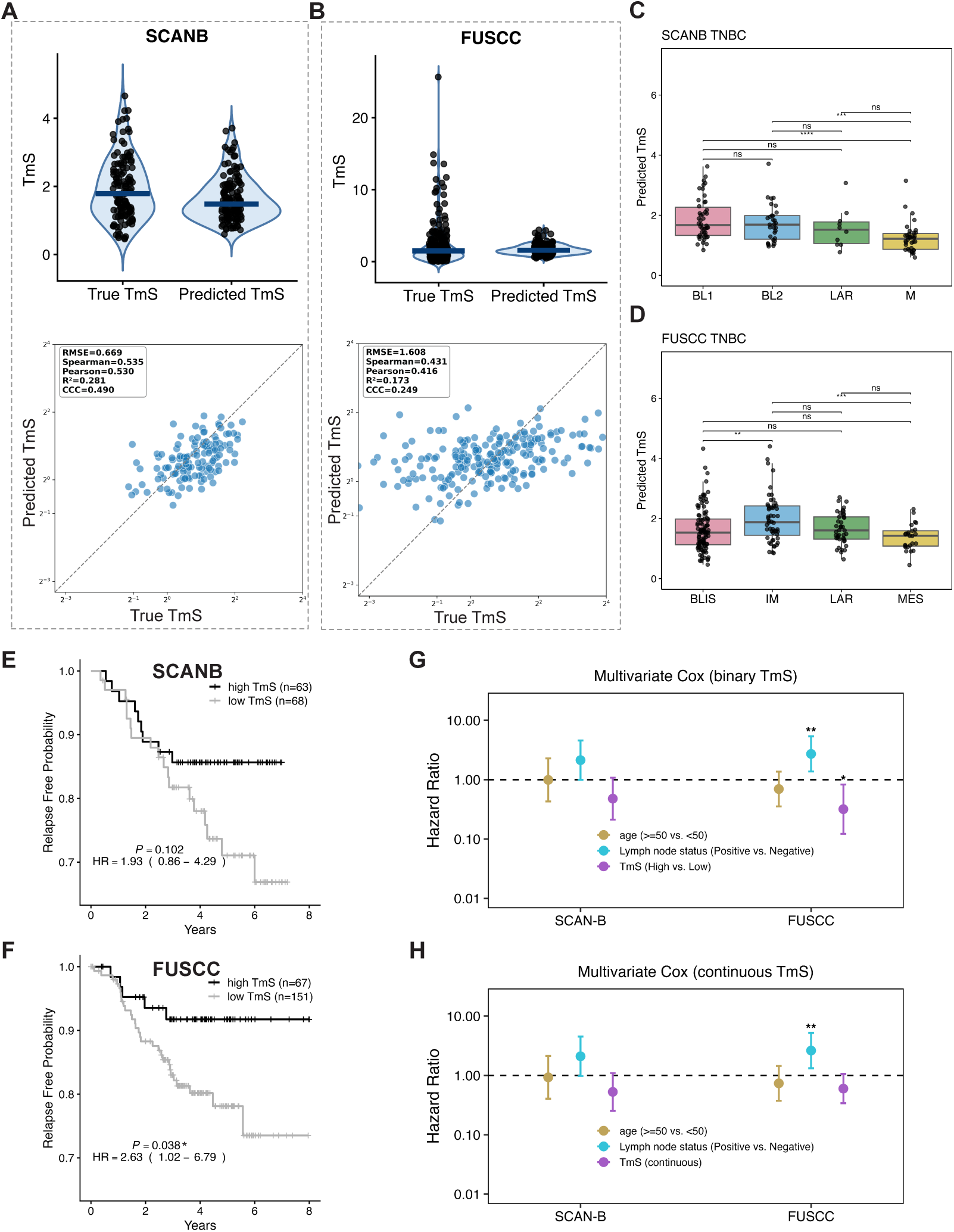
External Validation of TmSNet. Comparison of deconvolved (true) and predicted TmS values in the (**A**) SCAN-B and (**B**) FUSCC cohorts, shown using raw TmS distriubtions (top) and correlation metrics (bottom) (**C-D**) Predicted TmS values across established TNBC molecular subtypes in both external cohorts. (**E-F**) Relapse-free survival curves for stratified by high versus low predicted TmS values for (**E**) SCAN-B and (**F**) FUSCC cohorts. (**G-H**) Multivariable Cox proportional–hazard models assessing the prognostic value of TmS after adjustment for clinical covariates. ** p <0.01 *** p <0.001

To evaluate whether predicted TmS preserves known biological stratification, we examined its distribution across established TNBC molecular subtypes in the two cohorts. In SCAN-B, predicted TmS differed significantly across Lehmann subtypes (**Fig. 3C**, ANOVA p = 1.36 × 10⁻⁴). Basal-like 1 (BL1) tumors exhibited the highest median predicted TmS, followed by BL2 and LAR subtypes, while mesenchymal (M) tumors demonstrated lower transcriptional burden. A similar pattern was observed in FUSCC (**Fig. 3D**, ANOVA p = 1.22 × 10⁻³), indicating preservation of subtype-specific transcriptional gradients across institutions.

We evaluated the clinical association of the predicted TmS with recurrence-free survival and observed prognostic signal differed across cohorts (**Fig. 3E-H**). For SCAN-B, neither the binary comparison (log–rank p = 0.10; adjusted Cox p = 0.15; HR = 0.63, 95% CI 0.33–1.19) nor the continuous model (adjusted p = 0.16; HR = 0.69, 95% CI 0.41–1.16) is statistically significant, but the suggested trend is consistent with the reported association of high TmS with lower recurrence free survival. FUSCC shows a statistically significant Kaplan–Meier separation (log–rank p = 0.03), but the effect attenuates after adjustment (Cox p = 0.07; HR = 0.57, 95% CI 0.31–1.05) and is not statistically significant when modeled continuously (adjusted p = 0.20; HR = 0.78, 95% CI 0.52–1.15), still suggesting a trend toward lower risk in the high–TmS group.

## Discussion

Here, we present an orthogonal approach to the active research field of multi–omic deep learning–based patient subtyping by introducing tumor-specific total mRNA expression (TmS) as a biologically grounded biomarker to inform deep learning. Predicting clinical outcomes is challenging due to extensive molecular heterogeneity across patients, tumor regions, and time, which limits the reliability and generalizability of traditional biomarkers. Although multi–omic data can improve prediction, integrating these high–dimensional and noisy signals is difficult given tumor cell clonal diversity, stromal and immune admixture, and technical batch effects. TmS helps overcome these obstacles by capturing a core tumor property (i.e. cell-type specific total mRNA presence). As a lineage–agnostic, continuous metric, TmS summarizes complex molecular variation into an interpretable signal that enables deep learning models to more effectively link multi–omic patterns to clinical outcomes, supporting improved generalizability across cohorts. Finally, TmSNet proves promising to replace deconvolution-intensive procedures to predict TmS in one new patient at a time, ideal for clinical utility.

To contextualize TmSNet within the broader landscape of AI-driven cancer prediction, it is useful to compare this approach with deep learning models that use hematoxylin and eosin (H&E) whole-slide images as input. On two independent H&E datasets (lung and bladder), weakly supervised CNNs achieved moderate survival discrimination (C-index = 0.70 in lung; 0.61 in bladder) and remained independently prognostic after adjustment, suggesting that morphologic patterns alone can capture clinically relevant risk signals^41^. However, this achievement in prognosis prediction does not match up to the predictive performance of deconvolution-based TmS^13^. More recently, large-scale pathology foundation models such as CHIEF have advanced this field, demonstrating cross-institution generalization and survival prediction performance approaching a C-index of ∼0.74 across cancer types^42^. H&E-based deep learning has also been applied to molecular and therapeutic prediction tasks, including CNN-based prediction of HER2 status and trastuzumab response in breast cancer and transformer-based prediction of immunotherapy response and survival in non–small cell lung cancer^43,44^. However, image-based models remain sensitive to stain variability, scanner differences, and site-specific pathology workflows, which can limit generalizability and affect scaling up across institutions. In contrast, TmSNet predicts a biologically grounded, lineage-agnostic continuous biomarker (TmS) directly from multi-omic molecular profiles, providing complementary mechanistic insight when sequencing data are available.

Several recent frameworks have explored deep learning for multi-omic integration, including methods such as MOGONET, and DeepMOCCA, which aim to jointly model heterogeneous molecular layers to predict clinical outcomes or identify latent biological factors^26,27^. These approaches have demonstrated that integrating transcriptomic, epigenomic, and genomic features can improve predictive performance relative to single-modality models. However, many existing frameworks either rely on unsupervised latent factor discovery or directly predict categorical clinical endpoints such as survival groups or treatment response, which can obscure intermediate biological variation and limit interpretability. The performance of these models on a relatively small patient cohort of 2-300 is expected to be highly limited. In contrast, TmSNet adopts a supervised regression strategy centered on a biologically grounded intermediate phenotype. By focusing on predicting a continuous transcriptional output that reflects tumor purity, ploidy, and transcript abundance, the framework captures underlying molecular gradients associated with tumor aggressiveness while maintaining mechanistic interpretability. This intermediate-phenotype approach also reduces the dimensional complexity of the prediction task and enables the model to learn more stable representations of tumor biology across heterogeneous cohorts.

Extensive cross-validation across multiple cancer types demonstrated stable predictive performance across independent folds, and external validation in SCAN-B and FUSCC cohorts confirmed that the learned mapping generalizes beyond the TCGA training dataset. Although prediction error increased in external cohorts, common with external validation, it is likely due to the lack of DNA methylation data. More external validations will be performed as we continue to identify cohort datasets with matched input data with the selected models (**Table 1**). Alternatively, other accessible multi-omic input data to replace methylation array data will be explored. The clinical translation of TmSNet requires further prospective validation and standardization across cohorts and tumor stages before predictions can reliably inform real-world decision-making.

In summary, TmSNet provides a scalable and interpretable framework for predicting tumor transcriptome ploidy from multi-omic data^13^. By modeling a biologically grounded intermediate phenotype, the approach links heterogeneous molecular signals to clinically relevant tumor behavior. As multi-omic datasets continue to expand, frameworks such as TmSNet may help translate complex molecular measurements into robust biomarkers that improve patient stratification and therapeutic decision-making.

## Methods

### Dataset Description

#### Training cohorts

TmSNet was trained and internally validated using multi-omic data from 12 cancer types in The Cancer Genome Atlas (TCGA; total n = 3,699; cancer-type specific sample size ranges from 155 to 633). Raw data are available from the Genomic Data Commons Data Portal (https://portal.gdc.cancer.gov/). For each tumor sample, up to four data modalities were considered as candidate inputs: raw read counts from mRNA sequencing and miRNA sequencing, raw log ratio DNA methylation arrays, and immune cell-type proportions estimated by CIBERSORTx. The optimal feature combination for each cancer type was determined empirically via cross-validation and is reported, along with the sample size, in **Table 1**. Copy-number variation features were intentionally excluded to prevent information leakage, as tumor ploidy is a direct component of the TmS calculation. Deconvoluted TmS score, derived from matched bulk RNA and DNA sequencing, served as the ground truth labels for model training^13^.

#### External validation cohorts

We used two independent breast cohorts as the external validation cohorts. The Fudan University Shanghai Cancer Center (FUSCC)^39^ cohort (n= 218) and the Sweden Cancerome Analysis Network-Breast (SCANB)^40^ cohort (n= 131). Because DNA methylation data were unavailable in both external cohorts, only mRNA, miRNA, and CIBERSORTx were used as model inputs for external prediction.

The deconvolution TmS, which serves as the ground truth here, is expressed as^13^:

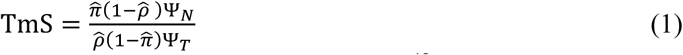

where 𝜋& denotes the tumor-specific mRNA proportion estimated using DeMixT^45^, Ψ_N_denotes ploidy of normal tissue, 𝜌& and Ψ_T_ denotes estimated tumor purity and ploidy estimated using ASCAT^46^, ABSOLUTE^47^, or Sequenza^48^ based on the matched DNA sequencing data, respectively. All TmS values are log_2,_transformed prior to model training and evaluation.

### TmSNet: A Multi-Omic Deep Learning Framework

We introduce TmSNet (Tumor mRNA Signal Network), a machine learning framework designed to predict total tumor mRNA (TmS) using multi-omic data. For each cancer type, TmSNet independently trains and evaluates multiple model configurations and selects the configuration with the highest cross-validated concordance correlation coefficient (CCC). The framework is described in detail by the following components: a. feature embedding, b. model training, c. model testing and validation.

### Feature embedding

For each cancer type, we implement an ensemble feature extraction strategy using three feature selection algorithms: extreme Gradient Boosting (XGB), LASSO regression, and Elastic Net (EN) for capturing nonlinear interactions and sparse, interpretable feature selection. Given input matrix 𝑋 ∈ ℝ^n×p^ where 𝑛 are number of samples and 𝑝 are number of features (e.g. gene names in mRNA profile) and the TmS output, selected features would be given by: 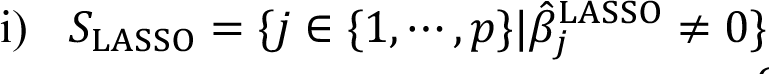, where 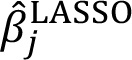 is the estimated coefficient for feature 𝑗 after solving LASSO, 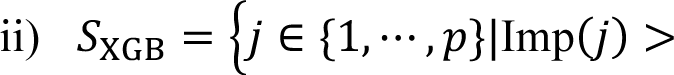 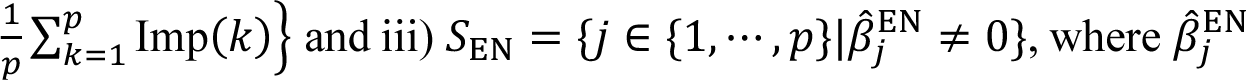 is the estimated coefficient for feature 𝑗 after solving EN. For each feature selection method, different feature combinations are trained independently producing their own feature subset selected 𝒮^m^ with |𝒮^m^| = 𝑑^m^ and 𝑚 is the indicator of a subset of the power set of mRNA, miRNA, methylation and CIBERSORTx.

### Model Training

The feature selection generates a reduced feature vector 𝒙_i,𝒮_ ∈ ℝ^d^ for sample *i*, a subset of the features used to train two separate regressor learners: a specialized encoder regressor and a feed-forward neural network (**Fig. 1B**). We denote 𝒙_i_ ≡ 𝒙_i,𝒮_ in the subsequent sections for simplicity.

### Specialized Encoder Neural Network (SENN) Training

FSENN is a specialized latent encoder regressor that is used to translate a feature vector 𝒙 into latent Gaussian space 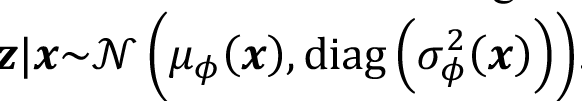. Reparametrizing of which gives us:

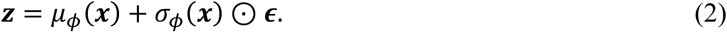

Here, ⊙ denotes the Hadamard product for element-wise multiplication between two vectors, and 𝝐 ∼ 𝒩(0, 𝐈) injects standard Gaussian randomness into the latent representation, enabling posterior sampling.

Given latent embeddings 𝒛, we proposed a regression head 𝑓_B_(⋅) for TmS prediction. We train the encoder parameters 𝝓 and regression-head parameters 𝜽 by minimizing the mean squared error:

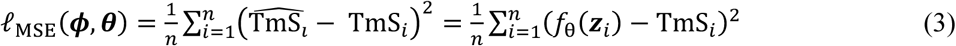

We compute gradients by backpropagation through the reparametrized latent 𝒛_=_ and the head 𝑓_B_(⋅). Parameters are optimized using Adam, with a single posterior sample drawn per forward pass^49^.

### Feed-Forward Neural Network (FFNN) Training

The FFNN design follows a standard multi-layer perceptron (MLP), with five hidden layers (𝐿 = 5). The architecture utilizes a hidden width of H = 98. We initialize inputs to the network as 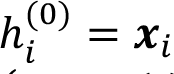. The first transformation applies the weights and a ReLU activation function as 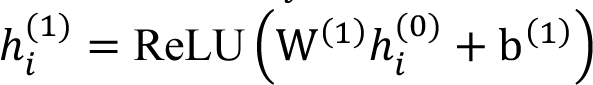; Subsequent hidden layers repeat the affine mapping and ReLU transform up to layer 5. The final prediction 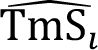 is generated by a linear transformation of the last hidden state: 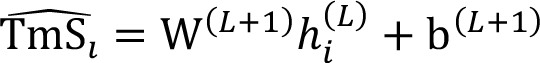, 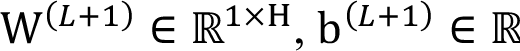.

The model is trained for 50 epochs by minimizing the mean squared error loss. Let 𝝎 denote the parameters of FFNN, the loss function is 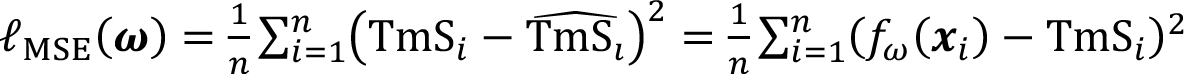.

### Hyperparameter Optimization

Given the SENN and FFNN architectures described above, hyperparameter configurations such as dropout rate and layer width remain to be determined. We employ Bayesian Optimization (BO) to search for the hyperparameter set 𝜆 within the search space Λ that minimizes the expected validation loss on the held-out set 𝑆_val_,

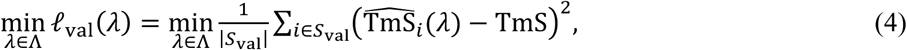

BO searches for potential hyperparameter configurations iteratively. At iteration 𝑡 − 1, the optimizer selects next hyperparameter configuration 𝜆_t_, based on existing dataset 𝒟_*t*_ = 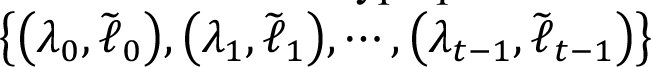. An acquisition function 𝛼_t_(𝜆) representing expected improvement balances exploration (via the uncertainty 𝜎_t_(𝜆) of 𝒟_t_) and exploitation (via the mean 𝜇_t_(𝜆) of 𝒟_t_) relative to the current best validation loss 𝓁̃^*^. The Gaussian CDF Ψ and PDF 𝜓 quantify the probability and magnitude of improvement, given by:

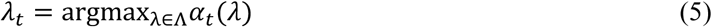

where 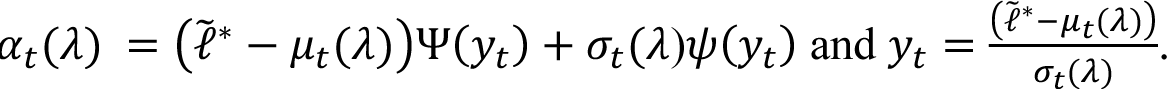

Post evaluation of the model at the chosen parameters 𝜆_t_, the updates of the surrogate model’s dataset by adding the new observation 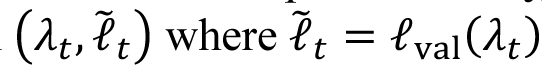. The updated dataset 𝒟_t+1_ refines the Gaussian-process surrogate, improving its prediction of the loss landscape for the next iteration is given by 𝓓_*t*+1_ = 𝓓_*t*_ ∪(λ_*t*_, 𝓁̃_*t*_).

### Data Imbalance Issue

We addressed data imbalance using Synthetic Minority Over-sampling Technique for Regression with Gaussian Noise (SMOGN)^50^. Given the all-numeric matrix with the designated regression target (TmS), SMOGN increases the effective sample density in target regions that are under-represented while preserving local covariate structure. For each rare instance, SMOGN locates k-nearest neighbors in feature space and interpolates between them to create new samples, with small Gaussian noise to preserve diversity. This maintains local feature–target structure while enriching under-represented outcome ranges.

### Model Testing and Validation

For external validation, the model is retrained on the entire TCGA training cohort using the best-performing configuration identified during cross-validation. The trained pipeline weights and scaler files are then applied to the external cohorts to generate TmS predictions. TmS predictions are generated from the selected features through either a latent-space encoding (SENN) or a direct feed-forward mapping (FFNN). Gene-expression matrices are standardized in the same way as the training step. Region-level TmSs are obtained for the TRACERX cohort while sample-level TmS are obtained for the CPTAC-PDAC, SCANB and FUSCC cohort. Tumors and tumor regions with purity outside (0.2,0.85) were excluded.

Predictive agreement between true TmS and predicted TmS is evaluated by RMSE, Pearson’s r(and R^2^), Spearman’s ρ_[_ and concordance correlation coefficient (CCC).

## Data Availability

TCGA multi-omic data used for model training are publicly available through the Genomic Data Commons Data Portal (https://portal.gdc.cancer.gov/).

FUSCC TNBC multi-omic data are accessible via the National Omics Data Encyclopedia (NODE: OEP000155; http://www.biosino.org/node).

Sequence data available in NCBI Gene Expression Omnibus (OncoScan array: GSE118527) and Sequence Read Archive (WES and RNA-seq: SRP157974). SCAN-B: somatic mutation data are available at https://data.mendeley.com/datasets/2mn4ctdpxp/1.

RNA-seq raw counts are available at https://data.mendeley.com/datasets/yzxtxn4nmd/3^52^. TmSNet model weights and analysis code will be made available on GitHub.

## Acknowledgments

The results presented here are in part based upon data generated by the TCGA Research Network: https://www.cancer.gov/tcga. The authors would like to acknowledge the SCAN-B study, all patients and clinicians participating in the SCAN-B study, personnel at the central SCAN-B laboratory at the Division of Oncology, Department of Clinical Sciences Lund, Lund University, as well as the Swedish national breast cancer quality registry (NKBC), Regional Cancer Center South (RCC Syd), Regional Biobank Center South (RBC Syd), and the South Sweden Breast Cancer Group (SSBCG). This work was supported by National Cancer Institute R01CA268380 (to W.W.), Department of Defense PC210079 (to W.W.). J.C.L is a TRIUMPH Fellow in the CPRIT Training Program (RP210028).

## Author Contributions

S.J, and W.W. conceived the project. A.P. developed the model architecture. A.P., J.C.L., S.J., C.F., Y.D., Y.D., performed the data analysis and interpretation. A.P., J.C.L, S.J., K.C., R.L., Q.T., M.M assisted with data pre-processing and data maintenance. A.P., J.C.L, S.J., A.S., S.K., W.W. provided expertise and advised on data interpretation. W.W. conceived the study, planned and supervised the work, performed the analysis, and wrote the manuscript collaborating with all the other authors. All authors contributed to the interpretation of the results and commented on and approved the final manuscript.

## Supplementary Figures

**Supplementary Table 1.** Correlation of candidate genes with TmS in breast cancer from TCGA. Shown is a curated list of genes responsible for regulating gene expression and respective TmS values, biological pathway identity, Pearson correlation coefficients, and p values.

| Gene | Functional category | Pearson r | P-value |
| --- | --- | --- | --- |
| *XPC* | DNA repair / genome maintenance | -0.51 | $< 2.2 \times 10^{-16}$ |
| *SETD2* | Chromatin regulation | -0.51 | $< 2.2 \times 10^{-16}$ |
| *SMARCA2* | Chromatin remodeling | -0.49 | $2.2 \times 10^{-16}$ |
| *NIPBL* | Chromosome cohesion / genome stability | -0.43 | $2.2 \times 10^{-16}$ |
| *RFC2* | DNA replication | 0.39 | $< 2.2 \times 10^{-16}$ |
| *CD40* | Immune signaling | 0.38 | $2.2 \times 10^{-16}$ |
| *POLR2A* | Transcription machinery | -0.38 | $2.2 \times 10^{-16}$ |
| *SF3B1* | RNA processing / splicing | -0.35 | $< 2.2 \times 10^{-16}$ |
| *FOXA1* | Transcription factor | -0.35 | $2.2 \times 10^{-17}$ |
| *RFC1* | DNA replication | -0.35 | $< 2.2 \times 10^{-16}$ |
| *MED12* | Transcription regulation | -0.34 | $2.2 \times 10^{-16}$ |
| *MAD2L2* | DNA repair / replication stress | 0.32 | $2.2 \times 10^{-16}$ |
| *RFC4* | DNA replication | 0.31 | $1.0 \times 10^{-14}$ |
| *WRN* | DNA repair / genome stability | -0.31 | $7.0 \times 10^{-15}$ |
| *KDM3A* | Chromatin regulation | -0.3 | $1.3 \times 10^{-14}$ |
| *PBX1* | Transcription factor | -0.28 | $9.8 \times 10^{-13}$ |
| *WEE1* | Cell cycle regulation | -0.28 | $4.3 \times 10^{-12}$ |
| *RNASE1* | RNA metabolism | 0.28 | $2.9 \times 10^{-12}$ |
| *PCNA* | DNA replication | 0.28 | $1.4 \times 10^{-12}$ |
| *POU5F1* | Stem cell transcription factor | 0.27 | $2.9 \times 10^{-11}$ |
| *CDK13* | Transcription regulation | -0.26 | $4.4 \times 10^{-10}$ |
| *SOX2* | Developmental transcription factor | 0.25 | $6.7 \times 10^{-10}$ |
| *CEBPA* | Transcription factor | 0.23 | $1.3 \times 10^{-8}$ |
| *POLD1* | DNA replication | 0.23 | $6.7 \times 10^{-9}$ |
| *PAX7* | Developmental transcription factor | 0.23 | $6.1 \times 10^{-9}$ |
| *POLA1* | DNA replication | -0.23 | $1.3 \times 10^{-8}$ |
| *DHX9* | RNA metabolism | -0.21 | $8.5 \times 10^{-8}$ |
| *GATA6* | Transcription factor | 0.2 | $1.2 \times 10^{-6}$ |
| *POLQ* | DNA repair | 0.19 | $1.3 \times 10^{-6}$ |
| *MYCN* | Oncogenic transcription factor | 0.18 | $4.4 \times 10^{-6}$ |
| *EEF2* | Translation | -0.18 | $4.7 \times 10^{-6}$ |
| *POLN* | DNA repair | -0.17 | $3.5 \times 10^{-5}$ |
| *TOP1* | DNA topology / replication | -0.17 | $4.0 \times 10^{-5}$ |
| *SMC1A* | Chromosome cohesion | -0.16 | $1.1 \times 10^{-4}$ |
| *SMARCA4* | Chromatin remodeling | 0.16 | $5.8 \times 10^{-5}$ |
| *FAS* | Immune signaling / apoptosis | 0.15 | $2.8 \times 10^{-4}$ |

**Supplementary Table 2.** Performance comparison of baseline and TmSNet models across cancer types. Predictive performance for TmS across five cancer types, reported as cross-validated coefficient of determination (R²), concordance correlation coefficient (CCC), and root mean squared error (RMSE). Baseline models use 5-fold cross-validated XGBoost trained on mRNA features only. TmSNet represents a multi-stage framework integrating feature selection (XGBoost, LASSO, or elastic net) with deep neural network architectures (specialized encoder neural networks [SENN] or feed-forward neural networks [FFNN]). Models are evaluated using mRNA alone or combined mRNA and DNA methylation (DNAm) inputs. Across all cancer types, TmSNet consistently improves predictive performance over baseline models, with further gains observed when incorporating multi-omic features.

| Cancer Type | R<sup>2</sup> | CCC | RMSE | Framework | Model | Input Features |
| --- | --- | --- | --- | --- | --- | --- |
| Breast cancer | 0.648 | 0.780 | 1.218 | Baseline | XGBoost | mRNA |
|  | 0.783 | 0.881 | 0.633 | TmSNet | XGBoost/SENN | mRNA |
|  | 0.796 | 0.889 | 0.627 | TmSNet | XGBoost/SENN | mRNA + DNAm |
| Lung adenocarcinoma | 0.475 | 0.657 | 1.330 | Baseline | XGBoost | mRNA |
|  | 0.596 | 0.750 | 0.699 | TmSNet | ElasticNet/FFNN | mRNA |
|  | 0.696 | 0.807 | 0.598 | TmSNet | ElasticNet/FFNN | mRNA + DNAm |
| Bladder urothelial carcinoma | 0.401 | 0.623 | 1.183 | Baseline | XGBoost | mRNA |
|  | 0.711 | 0.826 | 0.813 | TmSNet | ElasticNet/SENN | mRNA |
|  | 0.867 | 0.924 | 0.548 | TmSNet | ElasticNet/SENN | mRNA + DNAm |
| Kidney renal clear cell carcinoma | 0.545 | 0.694 | 1.316 | Baseline | XGBoost | mRNA |
|  | 0.680 | 0.807 | 0.670 | TmSNet | LASSO/SENN | mRNA |
|  | 0.796 | 0.883 | 0.533 | TmSNet | LASSO/SENN | mRNA + DNAm |
| Pancreatic adenocarcinoma | 0.437 | 0.603 | 1.664 | Baseline | XGBoost | mRNA |
|  | 0.466 | 0.601 | 0.946 | TmSNet | ElasticNet/FFNN | mRNA |
|  | 0.485 | 0.607 | 0.954 | TmSNet | ElasticNet/FFNN | mRNA + DNAm |

**Supplementary Table 3.** Performance comparison of baseline models across mult-omic inputs. Predictive performance for XGBoost across five cancer types, reported as cross-validated coefficient of determination (R²), concordance correlation coefficient (CCC), and root mean squared error (RMSE). Baseline models were trained using 5-fold cross-validation with XGBoost and included the following input features: mRNA alone, combined mRNA and DNA methylation (DNAm), or mRNA, DNAm, miRNA, and inflammatory cell ratios (CIBERSORTx).

| Cancer Type | R<sup>2</sup> | CCC | RMSE | Input Features |
| --- | --- | --- | --- | --- |
| Breast cancer | 0.648 | 0.780 | 1.218 | mRNA |
|  | 0.672 | 0.791 | 1.273 | mRNA + DNAm |
|  | 0.689 | 0.801 | 1.253 | mRNA + DNAm +<br>miRNA+ CIBERSORTx |
| Lung<br>adenocarcinoma | 0.475 | 0.657 | 1.330 | mRNA |
|  | 0.583 | 0.715 | 1.077 | mRNA + DNAm |
|  | 0.599 | 0.724 | 1.083 | mRNA + DNAm +<br>miRNA+ CIBERSORTx |
| Bladder<br>urothelial<br>carcinoma | 0.401 | 0.623 | 1.183 | mRNA |
|  | 0.425 | 0.636 | 1.162 | mRNA + DNAm |
|  | 0.395 | 0.633 | 1.139 | mRNA + DNAm +<br>miRNA+ CIBERSORTx |
| Kidney renal<br>clear cell<br>carcinoma | 0.545 | 0.694 | 1.316 | mRNA |
|  | 0.523 | 0.682 | 1.301 | mRNA + DNAm |
|  | 0.499 | 0.681 | 1.321 | mRNA + DNAm +<br>miRNA+ CIBERSORTx |
| Pancreatic<br>adenocarcinoma | 0.437 | 0.603 | 1.664 | mRNA |
|  | 0.490 | 0.629 | 1.564 | mRNA + DNAm |
|  | 0.485 | 0.628 | 1.574 | mRNA + DNAm +<br>miRNA+ CIBERSORTx |

**Supplementary Figure 1.**
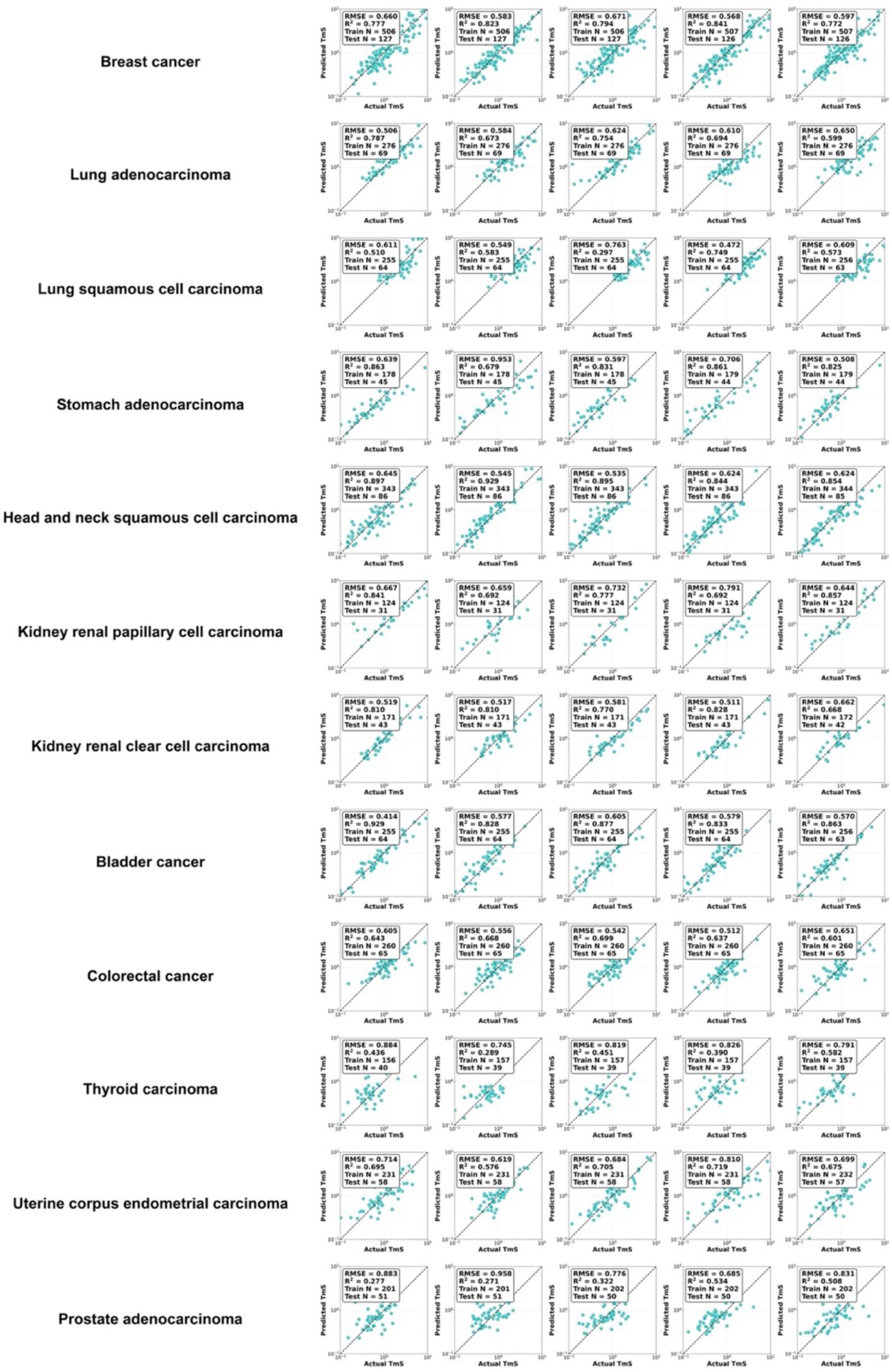
TmSNet predictive performance in the training TCGA cohort across different types of cancers with 5-fold cross-validation performance metrics RMSE and R^2^ across five folds. We performed 5–fold cross–validation performance of TmSNet across fourteen cancer types (rows), with each panel depicting predicted versus deconvolution–derived TmS on held–out samples from an individual fold. For every cancer type, the five panels correspond to the five validation folds; within each panel, points represent tumors from the test fold only, while the dashed identity line (y = x) provides a visual reference for perfect agreement. Each panel includes a performance inset reporting performance metrics RMSE and Pearson’s R².

**Supplementary Figure 2.**
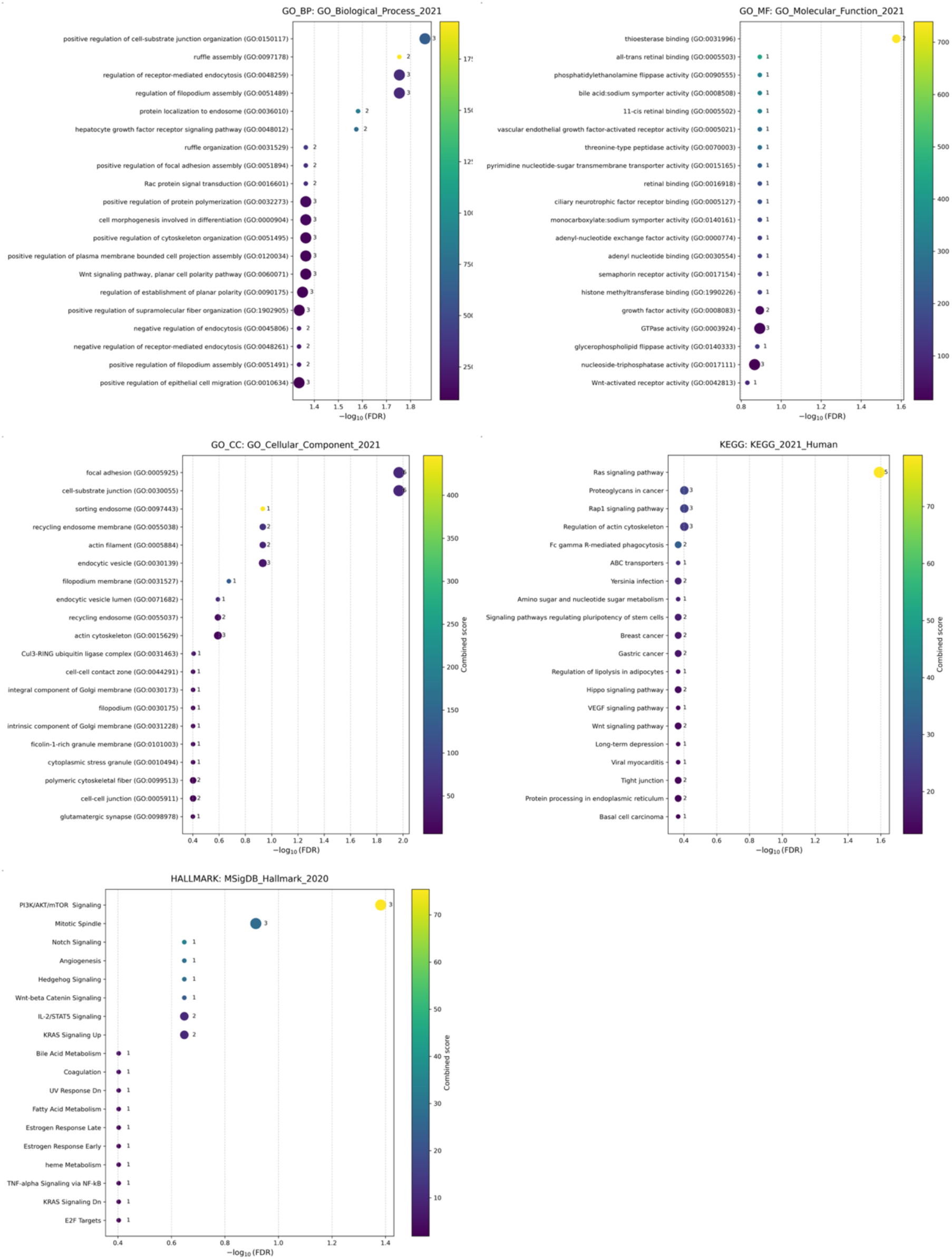
Pathway analysis from top features extracted by SHAP in three cancers from TCGA cohort using GO, Hallmark and KEGG modules for TCGA breast cancer.

**Supplementary Figure 3.**
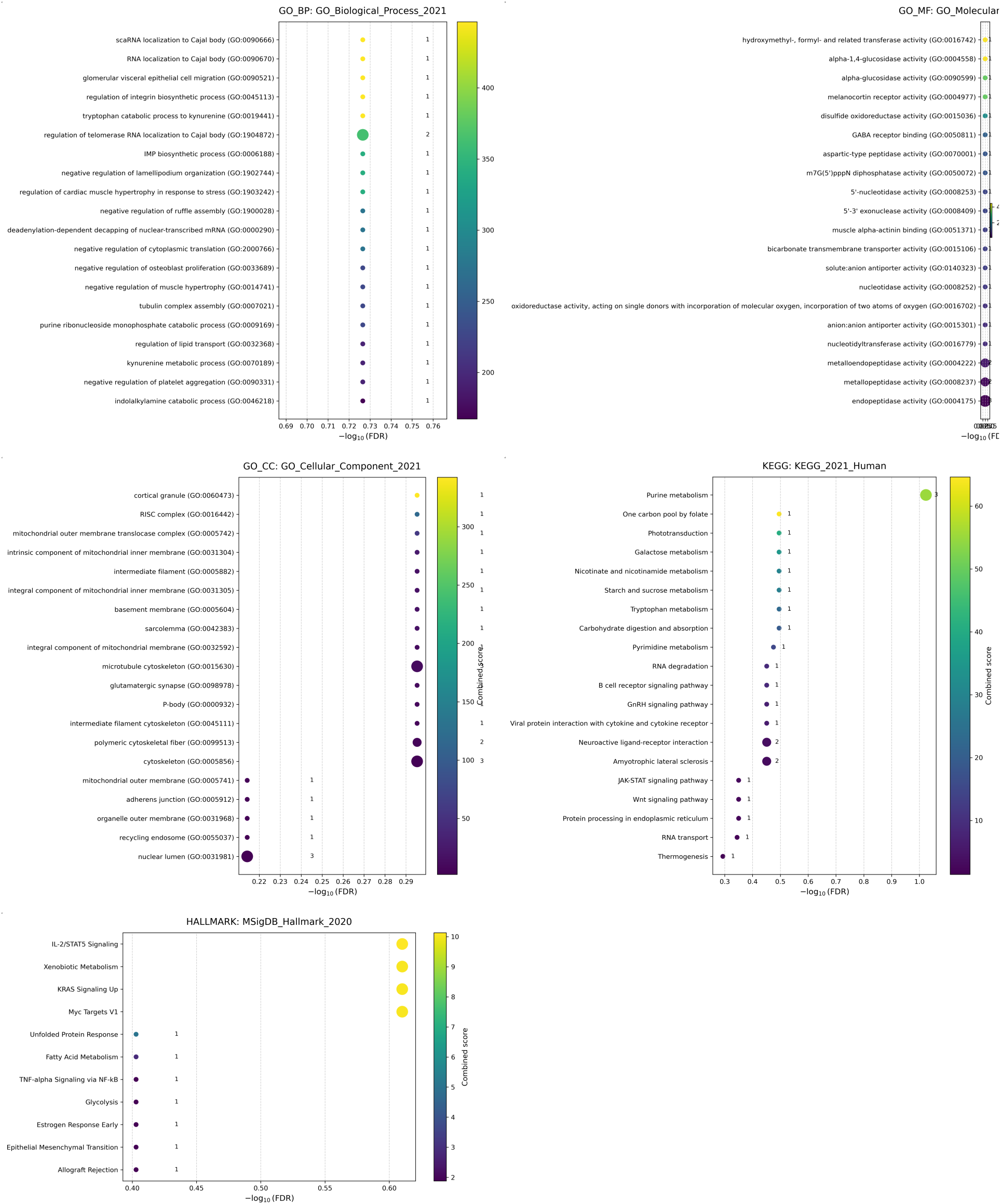
Pathway analysis from top features extracted by SHAP in three cancers from TCGA cohort using GO, Hallmark and KEGG modules for TCGA lung adenocarcinoma.

**Supplementary Figure 4.**
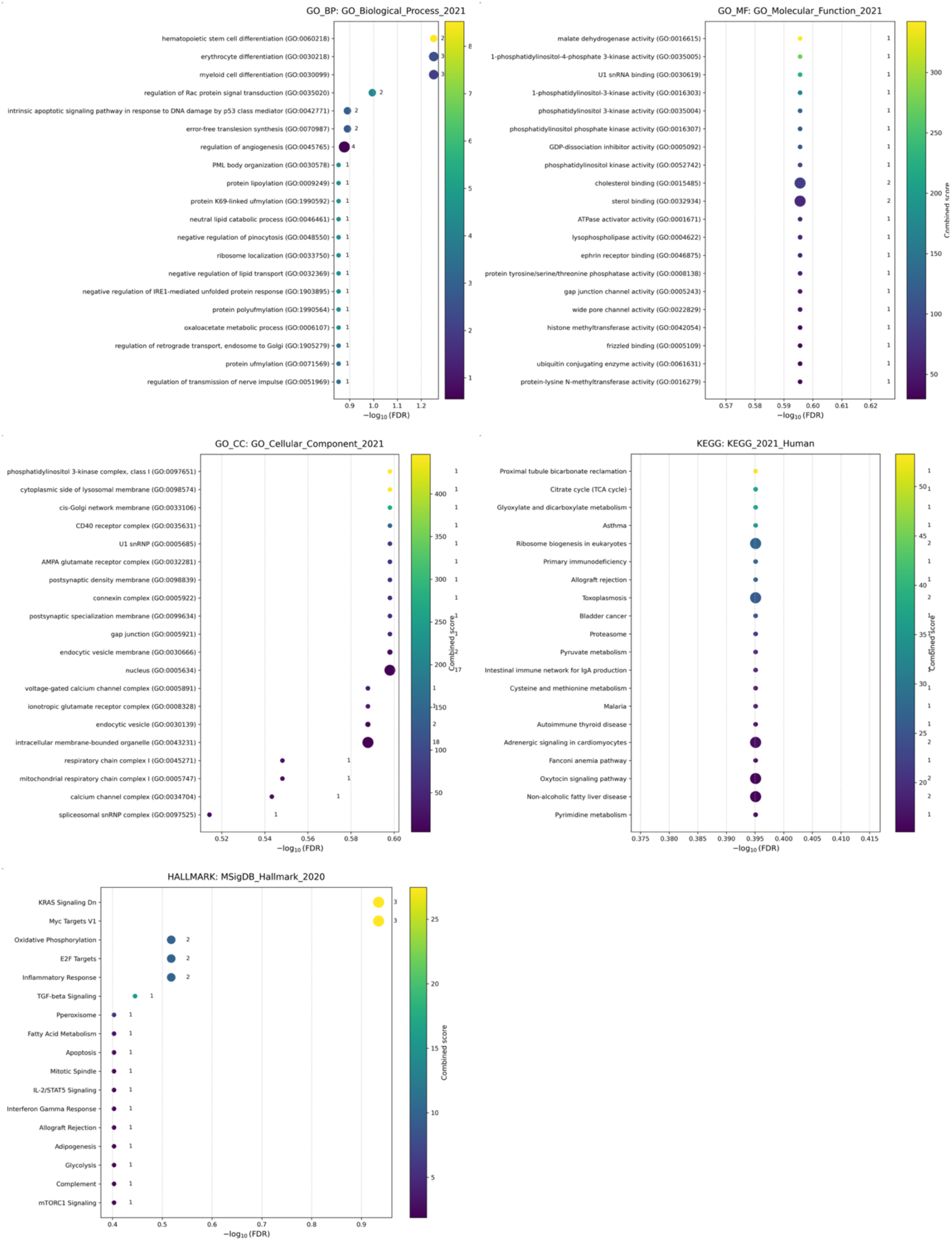
Pathway analysis from top features extracted by SHAP in three cancers from TCGA cohort using GO, Hallmark and KEGG modules for TCGA pancreatic cancer.

## References

1. Dentro, S. C. et al. Characterizing genetic intra-tumor heterogeneity across 2,658 human cancer genomes. Cell 184, 2239–2254.e39 (2021).

2. Quintanal-Villalonga, Á. et al. Lineage plasticity in cancer: a shared pathway of therapeutic resistance. Nat. Rev. Clin. Oncol. 17, 360–371 (2020).

3. Alizadeh, A. A. et al. Toward understanding and exploiting tumor heterogeneity. Nat. Med. 21, 846–853 (2015).

4. Mateo, J. et al. A framework to rank genomic alterations as targets for cancer precision medicine: the ESMO Scale for Clinical Actionability of molecular Targets (ESCAT). Ann. Oncol. 29, 1895–1902 (2018).

5. Li, M. M. et al. Standards and Guidelines for the Interpretation and Reporting of Sequence Variants in Cancer: A Joint Consensus Recommendation of the Association for Molecular Pathology, American Society of Clinical Oncology, and College of American Pathologists. J. Mol. Diagn. 19, 4–23 (2017).

6. Chakravarty, D. et al. Somatic Genomic Testing in Patients With Metastatic or Advanced Cancer: ASCO Provisional Clinical Opinion. J. Clin. Oncol. 40, 1231–1258 (2022).

7. Valpione, S. et al. Rechallenge with BRAF-directed treatment in metastatic melanoma: A multi-institutional retrospective study. Eur. J. Cancer 91, 116–124 (2018).

8. Chang, G.-C. et al. Predictive factors for EGFR-tyrosine kinase inhibitor retreatment in patients with EGFR-mutated non-small-cell lung cancer – A multicenter retrospective SEQUENCE study. Lung Cancer 104, 58–64 (2017).

9. Beroukhim, R. et al. The landscape of somatic copy-number alteration across human cancers. Nature 463, 899–905 (2010).

10. Dang, C. V. MYC on the Path to Cancer. Cell 149, 22–35 (2012).

11. Hanahan, D. & Weinberg, R. A. Hallmarks of Cancer: The Next Generation. Cell 144, 646–674 (2011).

12. Dai, Y. et al. Tumor microenvironment transcriptional activity enables robust stratification of chemotherapy response in triple-negative breast cancer. Cell Rep. Med. 7, 102610 (2026).

13. Cao, S. et al. Estimation of tumor cell total mRNA expression in 15 cancer types predicts disease progression. Nat. Biotechnol. 40, 1624–1633 (2022).

14. Ehteshami Bejnordi, B., et al. Diagnostic Assessment of Deep Learning Algorithms for Detection of Lymph Node Metastases in Women With Breast Cancer. JAMA 318, 2199–2210 (2017).

15. Liu, X. et al. A comparison of deep learning performance against health-care professionals in detecting diseases from medical imaging: a systematic review and meta-analysis. *Lancet Digit*. Health 1, e271–e297 (2019).

16. Goecks, J., Jalili, V., Heiser, L. M. & Gray, J. W. How Machine Learning Will Transform Biomedicine. Cell 181, 92–101 (2020).

17. Esteva, A. et al. A guide to deep learning in healthcare. Nat. Med. 25, 24–29 (2019).

18. Uyar, B. et al. Flexynesis: A deep learning toolkit for bulk multi-omics data integration for precision oncology and beyond. Nat. Commun. 16, 8261 (2025).

19. Ektefaie, Y. et al. Evaluating generalizability of artificial intelligence models for molecular datasets. *Nat*. Mach. Intell. 6, 1512–1524 (2024).

20. Geirhos, R. et al. Shortcut learning in deep neural networks. *Nat*. Mach. Intell. 2, 665–673 (2020).

21. Shi, L. et al. The MicroArray Quality Control (MAQC) project shows inter- and intraplatform reproducibility of gene expression measurements. Nat. Biotechnol. 24, 1151–1161 (2006).

22. Leek, J. T. et al. Tackling the widespread and critical impact of batch effects in high-throughput data. Nat. Rev. Genet. 11, 733–739 (2010).

23. Cabanillas Silva, P., et al. Longitudinal Model Shifts of Machine Learning–Based Clinical Risk Prediction Models: Evaluation Study of Multiple Use Cases Across Different Hospitals. J Med Internet Res 26, e51409 (2024).

24. Davis, S. E., Greevy, R. A., Lasko, T. A., Walsh, C. G. & Matheny, M. E. Detection of calibration drift in clinical prediction models to inform model updating. J. Biomed. Inform. 112, 103611 (2020).

25. Finlayson Samuel G., et al. The Clinician and Dataset Shift in Artificial Intelligence. N. Engl. J. Med. 385, 283–286 (2021).

26. Althubaiti, S., et al. DeepMOCCA: A pan-cancer prognostic model identifies personalized prognostic markers through graph attention and multi-omics data integration. *bioRxiv* 2021.03.02.433454 (2021) doi:10.1101/2021.03.02.433454.

27. Wang, T. et al. MOGONET integrates multi-omics data using graph convolutional networks allowing patient classification and biomarker identification. Nat. Commun. 12, 3445 (2021).

28. Wang, B. et al. Similarity network fusion for aggregating data types on a genomic scale. Nat. Methods 11, 333–337 (2014).

29. Ronen, J., Hayat, S. & Akalin, A. Evaluation of colorectal cancer subtypes and cell lines using deep learning. Life Sci. Alliance 2, e201900517 (2019).

30. Poirion, O. B., Jing, Z., Chaudhary, K., Huang, S. & Garmire, L. X. DeepProg: an ensemble of deep-learning and machine-learning models for prognosis prediction using multi-omics data. Genome Med. 13, 112 (2021).

31. Sharifi-Noghabi, H., Zolotareva, O., Collins, C. C. & Ester, M. MOLI: multi-omics late integration with deep neural networks for drug response prediction. Bioinformatics 35, i501–i509 (2019).

32. Mo, Q. et al. Pattern discovery and cancer gene identification in integrated cancer genomic data. Proc. Natl. Acad. Sci. 110, 4245–4250 (2013).

33. Kather, J. N. et al. Deep learning can predict microsatellite instability directly from histology in gastrointestinal cancer. Nat. Med. 25, 1054–1056 (2019).

34. Mobadersany, P., et al. Predicting cancer outcomes from histology and genomics using convolutional networks. Proc. Natl. Acad. Sci. 115, E2970–E2979 (2018).

35. Setty, M. et al. Characterization of cell fate probabilities in single-cell data with Palantir. Nat. Biotechnol. 37, 451–460 (2019).

36. Marjanovic, N. D. et al. Emergence of a High-Plasticity Cell State during Lung Cancer Evolution. Cancer Cell 38, 229–246.e13 (2020).

37. Newman, A. M. et al. Determining cell type abundance and expression from bulk tissues with digital cytometry. Nat. Biotechnol. 37, 773–782 (2019).

38. Lundberg, S. & Lee, S.-I. A Unified Approach to Interpreting Model Predictions. Preprint at 10.48550/arXiv.1705.07874 (2017).

39. Jiang, Y.-Z. et al. Genomic and Transcriptomic Landscape of Triple-Negative Breast Cancers: Subtypes and Treatment Strategies. Cancer Cell 35, 428–440.e5 (2019).

40. Saal, L. H. et al. The Sweden Cancerome Analysis Network - Breast (SCAN-B) Initiative: a large-scale multicenter infrastructure towards implementation of breast cancer genomic analyses in the clinical routine. Genome Med. 7, 20 (2015).

41. Qaiser, T. et al. Usability of deep learning and H&E images predict disease outcome-emerging tool to optimize clinical trials. *Npj Precis*. Oncol. 6, 37 (2022).

42. Wang, X. et al. A pathology foundation model for cancer diagnosis and prognosis prediction. Nature 634, 970–978 (2024).

43. Farahmand, S. et al. Deep learning trained on hematoxylin and eosin tumor region of Interest predicts HER2 status and trastuzumab treatment response in HER2+ breast cancer. Mod. Pathol. 35, 44–51 (2022).

44. Li, B. et al. Outcome-Supervised Deep Learning on Pathologic Whole Slide Images for Survival Prediction of Immunotherapy in Patients with Non–Small Cell Lung Cancer. Mod. Pathol. 36, 100208 (2023).

45. Wang, Z. et al. Transcriptome Deconvolution of Heterogeneous Tumor Samples with Immune Infiltration. iScience 9, 451–460 (2018).

46. Van Loo, P. et al. Allele-specific copy number analysis of tumors. Proc. Natl. Acad. Sci. 107, 16910–16915 (2010).

47. Carter, S. L. et al. Absolute quantification of somatic DNA alterations in human cancer. Nat. Biotechnol. 30, 413–421 (2012).

48. Favero, F. et al. Sequenza: allele-specific copy number and mutation profiles from tumor sequencing data. Ann. Oncol. 26, 64–70 (2015).

49. Alemi, A. A., Fischer, I., Dillon, J. V., & Murphy, K. Deep Variational Information Bottleneck. Preprint at http://arxiv.org/abs/1612.00410 (2019).

50. Branco, P., Torgo, L. & Ribeiro, R. P. SMOGN: a Pre-processing Approach for Imbalanced Regression. in Proceedings of the First International Workshop on Learning with Imbalanced Domains: Theory and Applications (eds Luís Torgo, P. B. & Moniz, N.) vol. 74 36–50 (PMLR, 2017).

51. Bergstra, J., Bardenet, R., Bengio, Y. & Kégl, B. Algorithms for Hyper-Parameter Optimization. in Advances in Neural Information Processing Systems vol. 24 (Curran Associates, Inc., 2011).

52. Staaf, J. et al. RNA sequencing-based single sample predictors of molecular subtype and risk of recurrence for clinical assessment of early-stage breast cancer. Npj Breast Cancer 8, 94 (2022).

